# The hypercubic Mk model in reduced state space for the coupled, reversible coevolution of multiple binary characters

**DOI:** 10.64898/2026.06.01.729317

**Authors:** Iain G. Johnston, Ramon Diaz-Uriarte, James D. Boyko

## Abstract

Many scientific questions involve the coevolution of coupled, binary features over time – from phenotypes in evolutionary biology to mutations in cancer development. Evolutionary accumulation models (EvAMs) often neglect reversibility in these systems, uncertainty in observations, and/or phylogenetic connections between observations. By contrast, the Mk model from phylogenetic comparative methods supports reversibility, uncertainty, and relatedness, but compute time scales like *O*(4^*L*^) in number of features *L*, making it challenging to apply to more than about six coupled, coevolving binary characters. Here, we introduce HyperMk2, a method using output from a Fitch-like parsimony algorithm to reduce the state space associated with many coevolving characters while retaining flexibility, reversibility, and phylogenetic information. This approach, while approximate, scales linearly in the number of distinct observations rather than exponentially in the number of characters, supporting the investigation of much larger systems than previously possible. We demonstrate how this method allows the inference of evolutionary dynamics of anti-microbial resistance in bacteria, including the identification of potential influences between characters, and discuss its broader application.

## Introduction

Many important processes in biology and disease involve systems acquiring (and losing) features over time. Learning about the dynamics by which these systems evolve can inform our basic scientific understanding – particularly about how these features might influence each other, rather than evolving independently – and also facilitate predictions of how new instances may evolve in future, and interventions to modify the evolution of the system.

Evolutionary accumulation modelling (EvAM) refers to the class of methods focused on studying the accumulation of binary features in evolutionary contexts (Diaz-Uriarte & Herrera-Nieto, 2022; Diaz-Uriarte & Johnston, 2025). EvAM, implicitly or explicitly, considers a state space of the 2^*L*^ possible different combinations of presence and absence for each of *L* different features, and the set of transition rates or probabilities between states. Acquisitions (and losses) are often pictured as occurring one-at-a-time, so the state space takes the form of a hypercube and “hypercubic inference” has been used to describe several EvAM approaches attempting to infer these transition rates from data (Aga et al., 2024; Johnston & Diaz-Uriarte, 2025; Moen & Johnston, 2023; Renz, Dauda, et al., 2025; Schill et al., 2020).

Evolutionary biology has long considered the evolution of characters on phylogenies, including ancestral state reconstruction and seeking evidence of coevolution (Boyko & Beaulieu, 2023; Pagel, 1997; Pagel & Meade, 2006; Revell & Harmon, 2022). These approaches include the Mk model for discrete trait evolution on phylogenies (Lewis, 2001; Pagel, 1994), a direct analogue of the Jukes-Cantor model originally developed for DNA sequence evolution (Jukes & Cantor, 1969). Methods here often assume reversible acquisition of features, and explicitly consider the phylogenetic coupling of observations; related work on states traversing a hypercubic state space is found in ontogenetic sequence analysis (Colbert & Rowe, 2008). But the number of coupled, coevolving characters that can be quantitatively considered together in these methods is highly limited (Johnston & Diaz-Uriarte, 2025).

A classic parallel example of EvAM is the study of cancer evolution, where a large field of methods has been developed to infer the ordering with which different mutations appear as a tumour develops (Beerenwinkel et al., 2015; Diaz-Uriarte, 2018; Diaz-Uriarte & Herrera-Nieto, 2022; Diaz-Uriarte & Johnston, 2025). Models often either seek to characterise deterministic dependencies between features (a tumour needs mutation *X*, or some combination of mutations, before mutation *Y* can appear) (Angaroni et al., 2022; Beerenwinkel et al., 2007; Desper et al., 1999; Gerstung et al., 2009; Montazeri et al., 2016; Nicol et al., 2021; Peterson & Kovyrshina, 2017; Szabo & Boucher, 2008), or influences in the form of changes to the rates of stochastic acquisitions (Aga et al., 2024; Greenbury et al., 2020; Hjelm et al., 2006; Schill et al., 2020). These models traditionally considered independent, cross-sectional, noiseless samples of the evolutionary process: samples from different patients reflecting independent realisations of the same underlying evolutionary process. More recently, methods have been developed considering the phylogenetic structure of tumour samples (Aga et al., 2024; Caravagna et al., 2018; Luo et al., 2023; Ross & Markowetz, 2016), and using a phylogenetic picture for inference of mutation ordering (Gao et al., 2022). Here, evolution is usually pictured as irreversible – once a mutation is acquired it is never lost, in a Dollo-like process (Gould, 1970) (although some approaches account for the fact that mutations may disappear in a tumour through intra-tumour competition). EvAM approaches handling phylogenetically-linked, uncertain data exist (Renz, Brun, et al., 2025), but reversibility has yet to be included in these methods.

The Mk (Markov k-state) model from evolutionary biology (Boyko & Beaulieu, 2021a; Lewis, 2001; Pagel, 1994) has been applied in EvAM circumstances (Johnston & Diaz-Uriarte, 2025). Here, a transition matrix (Q-matrix) describes transition rates between different states, and the likelihood of given observations is maximised over different parameterisations of this matrix. However, the core technology for inferring transitions in the Mk model, Felsenstein’s pruning algorithm (Felsenstein, 1973), has a time scaling of *O*(*n S*^2^), where *S* is the total number of states in the state space and *n* is the tree size. When considering *L* binary characters, the corresponding state space has 2^*L*^ states, and the corresponding *O*(*n* 4^*L*^) scaling makes anything more than simple systems (typically, *L* ≤ 7 (Johnston & Diaz-Uriarte, 2025)) intractable. Additionally, convergence of the parameter estimation process is challenged by the large number of parameters describing the transition rates between these states.

Overcoming this issue in an EvAM setting typically involves assuming irreversible dynamics and a strong model for ancestral state reconstruction invoking irreversibility and minimal evolution, which is applicable to some but certainly not all biological questions (Aga et al., 2024; Greenbury et al., 2020; Johnston & Williams, 2016; Moen & Johnston, 2023; Renz, Dauda, et al., 2025). A recent approach based on model selection provides a powerful reduction in search space – and selection of best model structures – for more general evolutionary problems (Boyko, 2026), but remains computationally challenging for systems containing many interacting binary characters.

In EvAM circumstances with cross-sectional data, we do not need to consider the full hypercubic space, and can readily consider a state space considering only the *K* observed unique binary strings (Giannakis et al., 2024; Vocht et al., 2026). For phylogenetically-coupled data, on the other hand, the ancestral states in the lineage leading to an observation must be considered, as the observation probability at each tip is conditional on ancestral states which act to couple observations. However, unless the evolutionary process covers a substantial fraction of the state space, some states will not provide useful or important contributions to the inference process. Our goal here is to use a parsimony picture to reduce the state space of the hypercubic Mk model to a tractable size for phylogenetically-embedded EvAM problems, while retaining the power of existing methods for Mk model fitting. We will further borrow from the EvAM philosophy and develop approaches to explicitly identify and test interactions between characters in evolution.

## Methods

### Reducing state space for the reversible hypercubic Mk model

We consider a Fitch-like (Fitch, 1971), partially greedy recursive algorithm for identifying sets of ancestral states that provide a parsimonious, yet flexible, state space for reversible accumulation modelling for a given system. We work starting from tip observations, first proposing a set of immediate ancestral states that are compatible with a minimum-evolution picture given their daughters. We then work toward the root of the tree, considering sets of ancestral states for each node that are minimum-evolution compatible with the sets in the daughter nodes. When the root is reached, each node has a collection of possible ancestral states that are minimum-evolution compatible with its daughters and sisters. The union of these sets is then used to create the reduced state space for the inference process, first by sampling possible states for each node, then constructing the set of transitions corresponding to each branch. This typically reduces the search space from 2^*L*^ to around 2*K* states, where *K* is the number of unique states.

The algorithm we use for this process relies on two operators applied to binary strings. The first operator is ⊕, applied to two individual binary strings, where *D*_1_ ⊕ *D*_2_ returns a set of binary strings *D* with discrepancies “expanded”. In other words, if *D*_1_ and *D*_2_ are identical at positions *a* and different at positions *b, D* will contain the set of binary strings that share *D*_1_ and *D*_2_‘s values at positions *a* and take all combinations of different values at positions *b*. For example, 001 *⊕* 101={001, 101} because *a*={2,3} and *b*={1}. In the case of uncertainty, we generalise so that any uncertain positions (in either string) are simply included in the set of discrepancy positions *b*. For example, 10? ⊕ 101={100, 101}.

The second operator is ⊗, applied to two sets (which may consist of single elements) of binary strings. This returns a tuple {*D, W*_1_, *W*_2_}. The first member *D* is the set of binary strings that minimise the summed Hamming Distance (HD) to an element of *D*_1_ and to an element of *D*_2_. In other words, we consider:

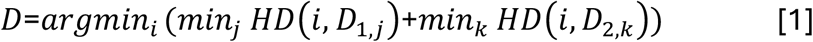

where *i* is an index over all binary strings of length *L*. Each of the two other returned members *W*_1_, *W*_2_ is a set of sets, containing “witnesses” for each of these minimum-HD values from each descendent node. Every element of *D* will have a corresponding set in *W*_1_ and a corresponding set in *W*_2_ (which may consist of single elements). From Eqn. 1, these give the sets of *j* and *K* strings in the two argument string sets corresponding to the minimum distance values. For example, {001,100} ⊗ {101} gives {*D*={001,100,101}, *W*_1_=D{001}, {100}, {001, 100}E, *W*_2_={{101}, {101}, {101}}. Here, the three states {001,100,101} all have a summed HD of 1 to an element of the two sets. First 001 is witnessed by 001 from the first argument and 101 in the second. Second, 100 is witnessed by 100 from the first argument and 101 in the second. Third, 101 is witnessed by either 001 or 100 from the first argument and 101 in the second.

The ⊕ operator can then be used to find a set of plausible ancestors for a pair of precisely-specified daughter observations. The ⊗ operator can be used to find a set of plausible minimum-evolution ancestor states for two descendant node sets, and to describe the subsets of descendant node states that correspond to these minimum-evolution pictures.

An algorithm for identifying a reduced state space for a given dataset is then given in Algorithm 1 and illustrated in Fig. 1, with a full example in Supp. Text 1. A computationally cheaper alternative, avoiding the storing of all witness-state relationships required for the recursive construction of a locally optimal tree, is given as Algorithm 2 in Supp. Text 2. Algorithm 1 in this article’s R implementation (not compiled code) took 15 minutes and used a maximum of around 3.5Gb of RAM for a problem with 22 binary characters and 118 observations; Algorithm 2 took seconds with negligible RAM usage (see Discussion). Algorithm 2, however, does not retain a “chain” of witness-state connections, and thus does not guarantee even locally that the minimum-evolution ancestor-descendant pairs are being chosen for a given branch. It instead samples randomly from the set of states constructed as minimum-evolution solutions. For this reason, and because even Algorithm 1 is likely to involve substantially less compute time than fitting the Mk model itself, we focus on Algorithm 1 in the illustrations and research here.

**Figure 1.**
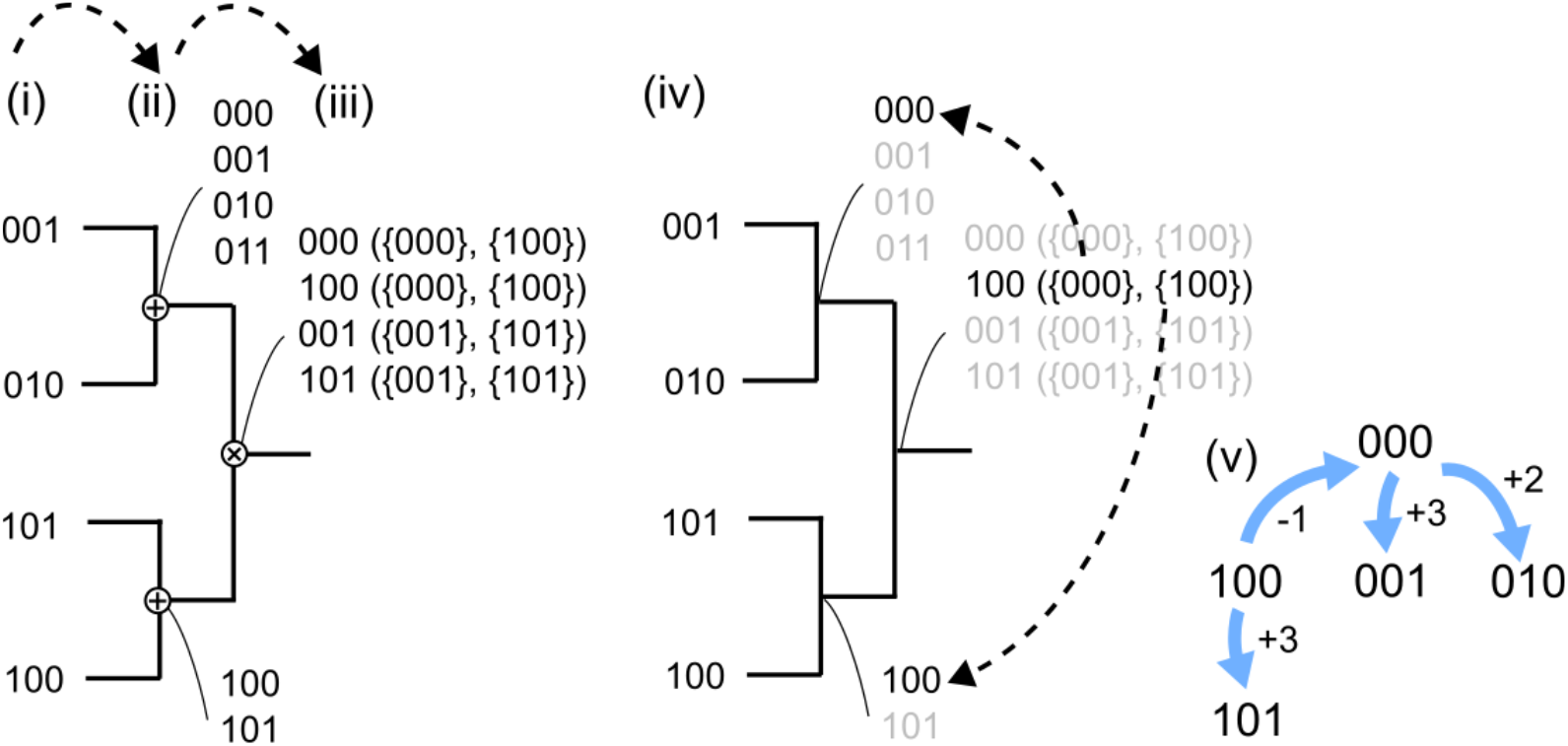
Illustration of state space reduction. Algorithms 1 and 2 begin with a set of binary string observations on a phylogeny (i). For each pair of observations, a set of minimal ancestral states is constructed (ii) by “expanding” every feature for which the descendant tips have different values using the ⊕ operator. (iii) Given two state sets for the descendants of an internal node, the ⊗ operator (see Methods) gives a set of minimum-evolution states of the node, and the “witnesses” for each of these states – the descendant states that correspond to each node state (in brackets). For example, in (iii), the possible ancestral state 000 is witnessed by the 000 state on the upper node and the 100 state on the lower node. The process continues towards the root until all nodes have been labelled with states and witnesses. (iv) A sampling approach then traverses the tree from the root towards the tips, choosing a particular state for the current node (in black) and reducing choices for the descendant nodes to witness sets for that state. Here, the choice of 100 for the root fixes all other nodes. (v) The corresponding reduced collection of states and gain/loss transitions between them, corresponding to inferred changes along branches in the tree. These state spaces will be drawn with the number of presence markers (1s) increasing down the figure. This pass through the algorithm is explicitly described in Supp. Text 1; alternative illustrations are shown in Supp. Fig. 1.

#### Algorithm 1

**Reducing state space for a hypercubic Mk model (Fitch-like)**

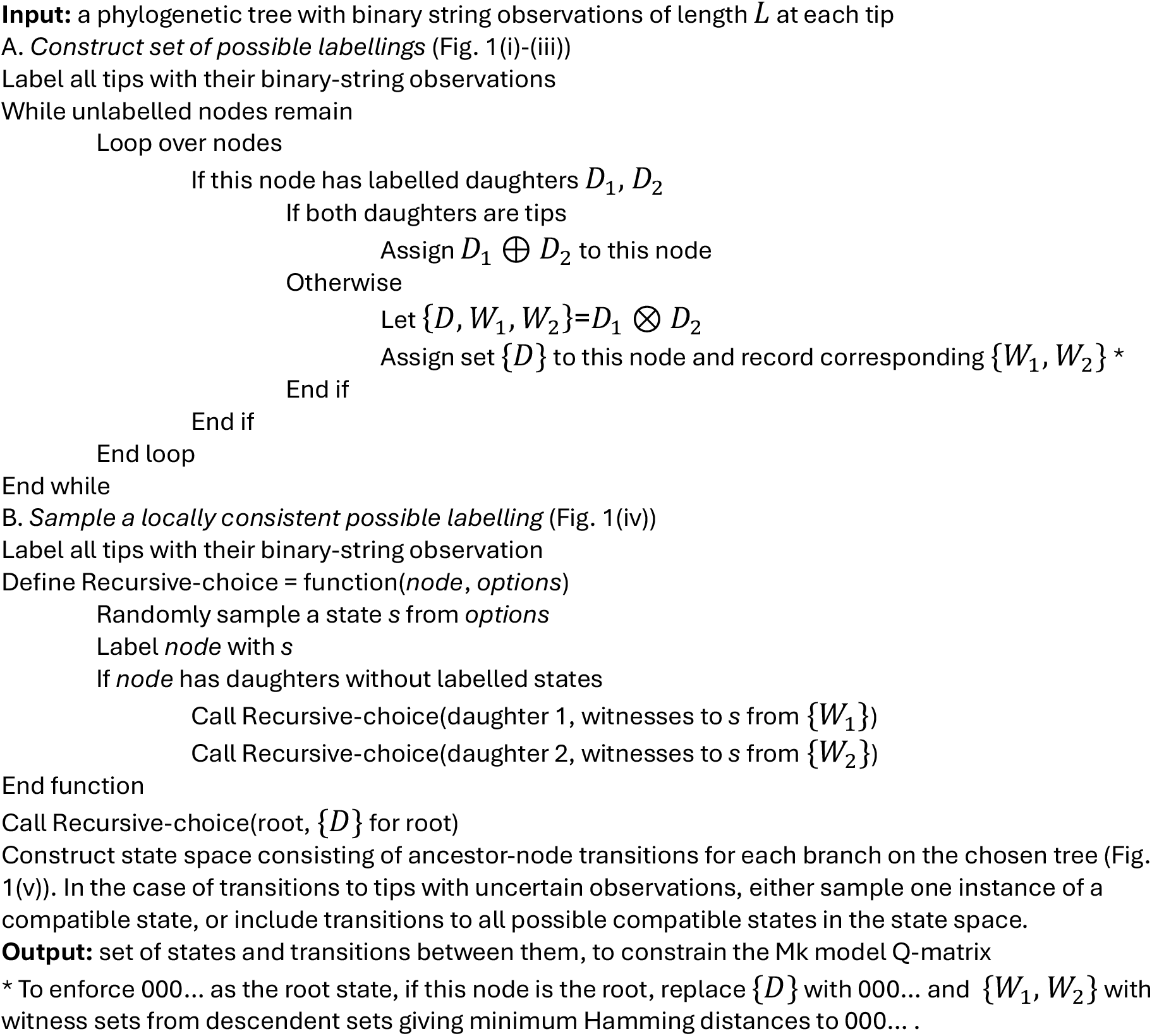

The ⊗ step only includes local information: there is no sense in which sets of states that might be useful or parsimonious for the EvAM model as a whole are considered more broadly than the sister pairs in this calculation. Doing so would generally involve a combinatoric search over the tree, which is similar to a demonstrably NP-hard problem even when reversibility is not considered (Giannakis et al., 2024). We proceed acknowledging that the reduced state space that this approach provides is not necessarily globally optimal from the minimal-evolution perspective. We also note that the aim of this approach is not to find a best estimate for the true ancestral states and trajectories (Boyko et al., 2026), but to identify a set of states that can parsimoniously explain the observed tip state.

In several EvAM contexts, the root state is known independently to be 000…, meaning the absence of any features. In this case, the (*) change can be made to Algorithm 1, to enforce the choice of this state for the root and support sampling of descendent states compatible with this state.

### Reduced state space for an irreversible hypercubic Mk model

We also consider the case of irreversible evolution, where transitions are allowed from absence to presence (0 to 1) but not vice versa (1 to 0). In this case, we can use our recent method adapting an algorithm from Gutin (Bang-Jensen & Gutin, 2018; Gutin et al., 2008) to the hypercubic transition graphs in EvAM (Giannakis et al., 2024). This algorithm produces a directed tree (an arborescence) in the binary state space that reaches every observation in a dataset with a minimum amount of “out-branching”. This produces a tree that involves the lowest and latest possible out-degrees for each space (corresponding to minimising evolutionary “choice”, particularly at earlier evolutionary times). The states in this parsimonious tree give a natural choice of reduced state space for problems involving irreversible evolution.

### Fitting the hypercubic Mk model in a reduced state space

Once the state space is established, we uniquely label each state and specify an all-rates-different (ARD) model for each transition in the set returned by Algorithm 1 (or 2) (Supp. Text 1). Only those transitions that appear in this output are allowed to take nonzero values in the Q-matrix describing the model, so both state and transition counts are reduced compared to a hypercubic model (Johnston & Diaz-Uriarte, 2025). We fit the resulting Mk model, specified by the original tree, tip states, and reduced Q-matrix, using the *castor* R package (Louca & Doebeli, 2018). Depending on biological context, we either enforce that the root node has state 000 … (in the case of accumulation proceeding from a known ancestor with no characters present) or allow a uniform prior over root node states.

In the case of tips with uncertain observations, there is a choice of protocol. We can include uncertainty over tip states naturally in the Mk model fit by assigning uniform probabilities over all compatible states (Johnston & Diaz-Uriarte, 2025; Louca & Doebeli, 2018). Or we can choose a specific compatible state for each tip with uncertain observations, chosen differently for each sample (Supp. Fig. 1B).

Each transition can in general involve the acquisition and loss of multiple characters simultaneously. In several EvAM use cases, this is a natural picture: for example, in the evolution of bacterial AMR, where multiple AMR characters can be gained and lost simultaneously via (for example) plasmid gain and loss. In other cases, these multi-change transitions reflect sets of transitions for which distinctions are not identifiable given the data and state space. This does not imply that all transitions must occur at the same time point, only that we cannot choose between the possible orderings of events for that set of transitions.

### Identifying interactions and departures from the null model of independent character evolution

The default null model for character evolution is that of independent characters, gained and lost with rates independent of any other characters in the system. This picture can readily be characterised within the HyperMk2 framework, simply by running the Mk model inference for each character independently, and summing the resulting log-likelihoods and AICs (for example). In this way, each character’s gains and losses are fit independent of other character behaviours, and the capacity of this null model to explain multi-characters observations is quantified. When considering evidence for interactions between features X and Y, we compare the total AIC of this null model for independent X and Y with the AIC of HyperMk (without any search space reduction) for X and Y together (i.e. an all-rates-different model involving states 00, 01, 10, and 11) (Johnston & Diaz-Uriarte, 2025). A large ΔAIC penalty for the independent model suggests support for pairwise influence between X and Y. This pairwise approach does not consider summed contributions from all features simultaneously as in other EvAM methods (Aga et al., 2024; Hjelm et al., 2006; Schill et al., 2020), but estimates the magnitude and direction of correlations between individual feature pairs (Grundler, 2025). While other null model structures can be considered to reduce potential bias (Boyko & Beaulieu, 2023), we implement this here for simplicity of interpretation.

### Visualisation and comparison of inferred transition networks

We will use a previously introduced visualisation for transition networks (Aga et al., 2024), where states are arranged in vertical layers by number of acquired features, and transitions between states are represented as edges (Fig. 1(v)). We will consider the probability “flux” through transitions, corresponding to the proportion of random walks on the transition network that traverse each edge. By contrast with the raw probability associated with a transition, this flux accounts for the probability of having reached the source state. To allow comparisons across approaches that may or may not support reversibility and/or multiple feature changes in a transition, we will use a “relative ordering matrix” 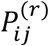, giving the probability that feature *j* is already present when feature *i* is acquired (analogous to the matrices used in (Dauda et al., 2025). This quantity is well defined for all EvAM approaches and serves as a quantitative comparison of stochastic dynamics.

### Implementation

Code for an R package implementing this approach is freely available at https://github.com/StochasticBiology/hypermk2; the set of key functions is described in the package documentation and in Supp. Table 1. For plotting and followup analysis, this has been embedded in the hypercubic inference ecosystem at https://github.com/StochasticBiology/hyperinf. The R libraries we use most directly are *castor* (Louca & Doebeli, 2018) for Mk model fitting, *ape* (Paradis & Schliep, 2019), *phangorn* (Schliep, 2011) and *phytools* (Revell, 2024) for phylogenetic operations, *igraph* (Csardi & Nepusz, 2006) and *stringr* (Wickham, 2019) for manipulating strings and graphs, and *ggplot2* (Wickham, 2011, 2016), *ggpubr* (Kassambara, 2020), and *ggraph* (Pedersen, 2020) for visualisation.

## Results

### The hypercubic Mk model in a reduced state space

We call our approach HyperMk2, for the second instance of our hypercubic Mk model picture (Johnston & Diaz-Uriarte, 2025). The algorithm for producing a reduced state space is described in the Methods and illustrated in Fig. 1. Briefly, we exploit the hypercubic nature of the state space in EvAM problems along with a picture of evolutionary parsimony to dramatically reduce the number of states we consider. We work backwards in time, first constructing sets of plausible minimal-evolution states for the immediate ancestors of observed tips. We then randomly sample states from these sets, iteratively reconciling them across sisters and to earlier node ancestors. This typically gives a set of states of size *O*(2*n*) rather than *O*(2^*L*^), with a number of transitions linear in *n*, which in turn constrains the form of the Q-matrix to be used in Mk model fitting. Different samplings will generally produce different state and transition sets; in the following we explore the consistency of the output across random samples. We fit an Mk model (using the *castor* R package (Louca & Doebeli, 2018)) to this reduced state space and interpret the inferred transition rates through an EvAM lens (Johnston & Diaz-Uriarte, 2025).

We must immediately note that the transitions in the reduced state space are not constrained to be single-step (in the sense of exactly one gain or loss of a feature). Transitions through the reduced state space may in general involve simultaneous gains and losses of multiple features – not necessarily meaning that this reflects biological reality, but that a parsimonious description of the dynamics cannot identify different subpathways within this set (Supp. Fig. 1A). The potential for multiple changes, and losses of features, makes several traditional ways of visualising and interpreting EvAM outputs (for example, the probability that a given feature is acquired at a given “step” in the evolutionary process) less meaningful. For this reason, we focus on two modes of output: the weighted transition networks themselves, and ordering matrices, giving the probability that feature A is present when feature B is acquired. These objects are quantitatively comparable across all EvAM methods (Aga et al., 2024; Dauda et al., 2025) and are directly tied to interpretations and predictions of the underlying dynamics (Johnston, 2026).

### Synthetic data test sets

We first test the performance of HyperMk2 on EvAM problems where the underlying generative dynamics are known. To this end we simulated a common EvAM test situation (Giannakis et al., 2024): two mutually repressing evolutionary pathways, one involving acquisition of feature 1, then feature 2, and so on, and the other involving acquisition of feature *L*, then *L*-1, and so on. We simulated this accumulation process on a randomly generated phylogeny (from a birth-death process (Paradis & Schliep, 2019)), giving observations in Fig. 2A. Such a system, involving two mutually exclusive evolutionary pathways, is observed in (for example) the reductive evolution of mitochondria across eukaryotes (Glastad & Johnston, 2025) and requires positive and negative interactions coupling features to realise, serving as a test case for these aspects of inference (Aga et al., 2024).

**Figure 2.**
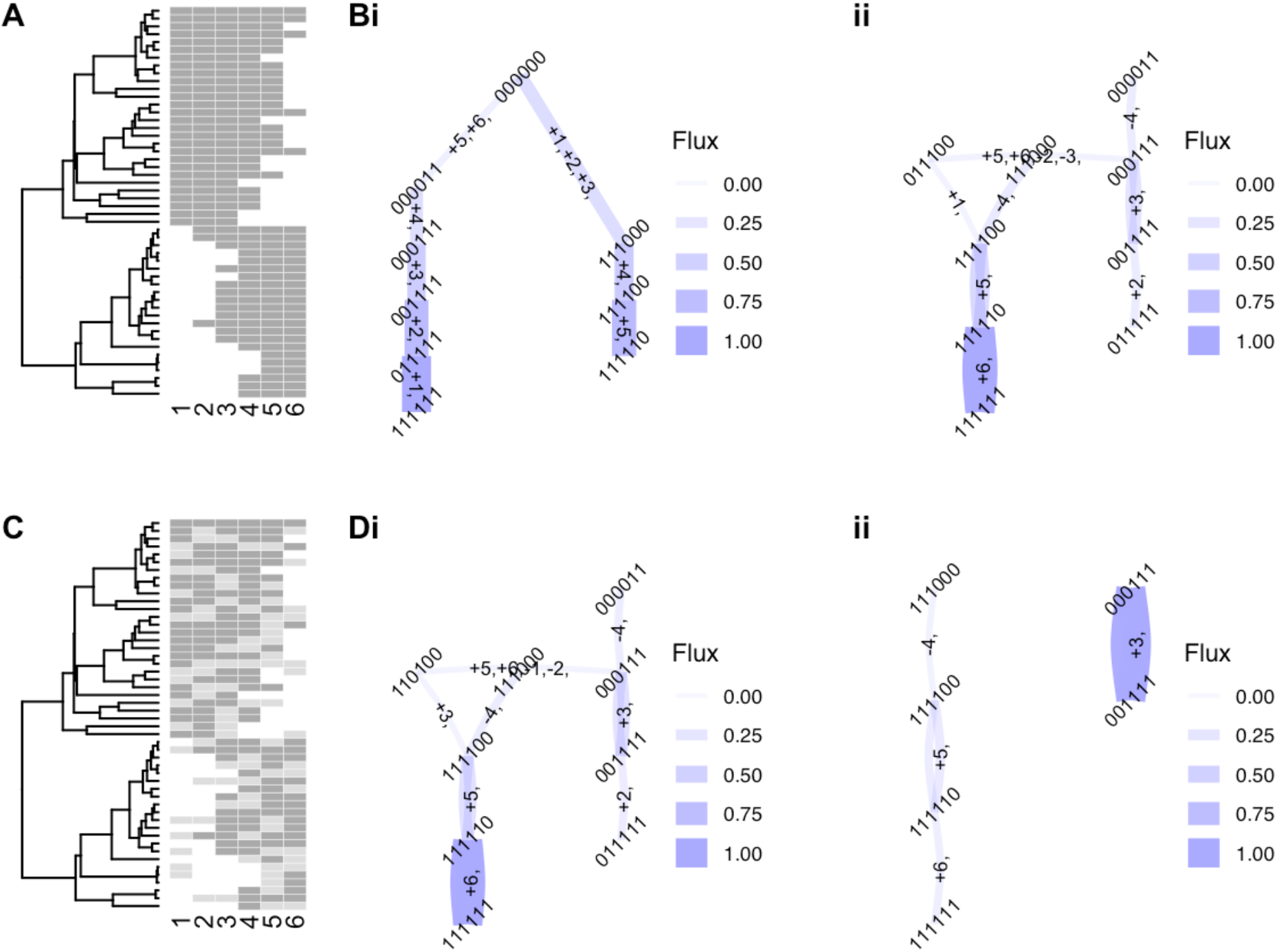
HyperMk2 inference for competing evolutionary pathways. **(A)** Synthetic data, where dark grey corresponds to presence and white corresponds to absence for each of 6 features. Accumulation progresses either in order 1-2-3… or order 6-5-4… on a randomly simulated tree (from the birth-death process). **(B)** Most common transitions from the inferred transition network from the (i) irreversible and (ii) reversible HyperMk2 model. As in Fig. 1iv, states are vertically ordered by their number of presence markers. Edges correspond to transitions, with width corresponding to the flux of random walkers simulated on the network through each. Fluxes below a threshold (0.05) are omitted from the figure. **(C)** Synthetic data with 50% artificially occluded (light grey corresponds to an uncertain marker). **(D)** Transition networks inferred from the reversible HyperMk2 model with (i) 10% and (ii) 40% of the data artificially occluded. In (B), (C), and (D)(i) the two mutually repressing evolutionary pathways are clearly inferred; their presence is still visible in (D)(ii). Supp. Fig. 2 shows the collection of inferred transition networks and dynamic summaries across samplings from the algorithm; Supp. Fig. 3 shows outputs for a wider range of occlusion proportions.

In one illustrative sampling, the irreversible picture of HyperMk2 recovered the transitions in Fig. 2B; the reversible picture recovered the transitions in Fig. 2C. These transitions are reported in terms of probability fluxes: the proportion of random walkers on the transition network that traverse that transition (see Methods). In both cases, the structure of the competing evolutionary pathways is naturally recovered, following results from other EvAM approaches (Aga et al., 2024; Greenbury et al., 2020; Johnston & Diaz-Uriarte, 2025; Moen & Johnston, 2023). Repeating the process across many random samplings for the reduced state space gave consistent results for the inferred dynamics (Supp. Fig. 2).

This test case also immediately demonstrates the ability of EvAM methods to reveal coevolutionary dynamics beyond the “null model” case of independently evolving characters. This two-pathway system clearly cannot be realised with independent characters: the acquisition of later features is completely dependent on the previous acquisition of earlier ones. Correspondingly, the AIC of the HyperMk2 model gives a clear favouring over that of the null model of independent character evolution (122.4 vs 181.1 in this example).

HyperMk2 can also handle uncertainty in observed data, expanding uncertain positions over all compatible values in the same way as discrepant positions are expanded over by the ⊕ operator, and fitting the Mk model either with a collection of sampled states or a probability distribution over possible states (Johnston & Diaz-Uriarte, 2025; Louca & Doebeli, 2018) (see Methods and Supp. Fig. 1B). To illustrate this capacity, we artificially occluded different proportions of the bilinear synthetic data, replacing observations with uncertainty markers (Fig. 2C). Fig. 2D shows that HyperMk2 recovers the generative evolutionary pathways for this case study readily for 10% occlusion and to some extent for 40% (Supp. Fig. 3 illustrates networks and ordering matrices as the proportion of occluded data is increased).

To demonstrate compatibility with previous approaches, we also simulated a simple single-pathway system in a smaller case study facilitating comparison with the previous HyperMk approach considering the full hypercubic state space of 2^*L*^ states (Johnston & Diaz-Uriarte, 2025). As shown in Supp. Fig. 4A-C, this approach readily matches the inferences from that more general picture in both the reversible and irreversible cases; in fact providing a more parsimonious picture of the underlying dynamics in this specific instance. Again, outputs across different samplings of the reduced state space gave consistent results (Supp. Fig. 4D-E).

### Antimicrobial resistance evolution

We next explore an evolutionary question that is both of wide societal impact (Baquero et al., 2021; Naghavi et al., 2024) and an expanding topic within EvAM (Renz, Dauda, et al., 2025) – the evolution of anti-microbial resistance (AMR) in bacteria. Bacteria acquire AMR by gaining genes conferring resistance, either through *de novo* mutations or the acquisition of genes via horizontal gene transfer (Dadgostar, 2019; Holt et al., 2015).

Chromosomal mutations may be lost in the absence of selective pressure applied through drug presence, and (for example) AMR-conferring plasmids may readily be lost in such conditions, as they incur a cost for carriage and maintenance. While some useful information about such reversible processes can be established from EvAM methods assuming irreversibility (Johnston, 2026), a framework naturally allowing for reversibility is a more robust and aligned alternative (Renz, Dauda, et al., 2025).

To explore the use of HyperMk2 in AMR data, we first looked at a classical dataset on multi-drug resistance in *Mycobacterium tuberculosis* (TB) (Casali et al., 2014), used to benchmark several EvAM approaches in the past (Aga et al., 2024; Greenbury et al., 2020; Johnston & Diaz-Uriarte, 2025; Moen & Johnston, 2023). Here, genomic data is used to infer resistance patterns against 10 drugs in TB isolates from Russia (a subset of data shown in Fig. 3A) and to infer the phylogenetic relationship between isolates. We applied HyperMk2 to this sample, resulting in state spaces with around 25 states and 35 transitions (compared to 1024 states in a full hypercubic model, and 5120 transitions (or 10240 for a reversible model), and 100 parameters in a pairwise-interaction (mutual hazards) model). The reduced state spaces, and inferred stochastic dynamics on them, were largely consistent across samplings of the algorithm (Fig. 3B). The relative orderings of feature acquisitions, with *INH, RIF*, and *STR* preceding all others with high probability, *EMB* acquired early, *PRO* and *PZA* later, and *AMI* late, are consistent with known behaviour and with the outputs of other EvAM approaches neglecting reversibility (Fig. 3C). These other approaches, HyperTraPS (Aga et al., 2024) and HyperHMM (Moen & Johnston, 2023), use a very strong assumption for ancestral state reconstruction in an irreversible picture – each ancestral state is given by the AND operator applied to descendant states. Then, an ancestor is assumed to have a feature if and only if both descendants have that feature – reflecting rare, irreversible acquisitions. HyperMk2 relaxes this picture substantially.

**Figure 3.**
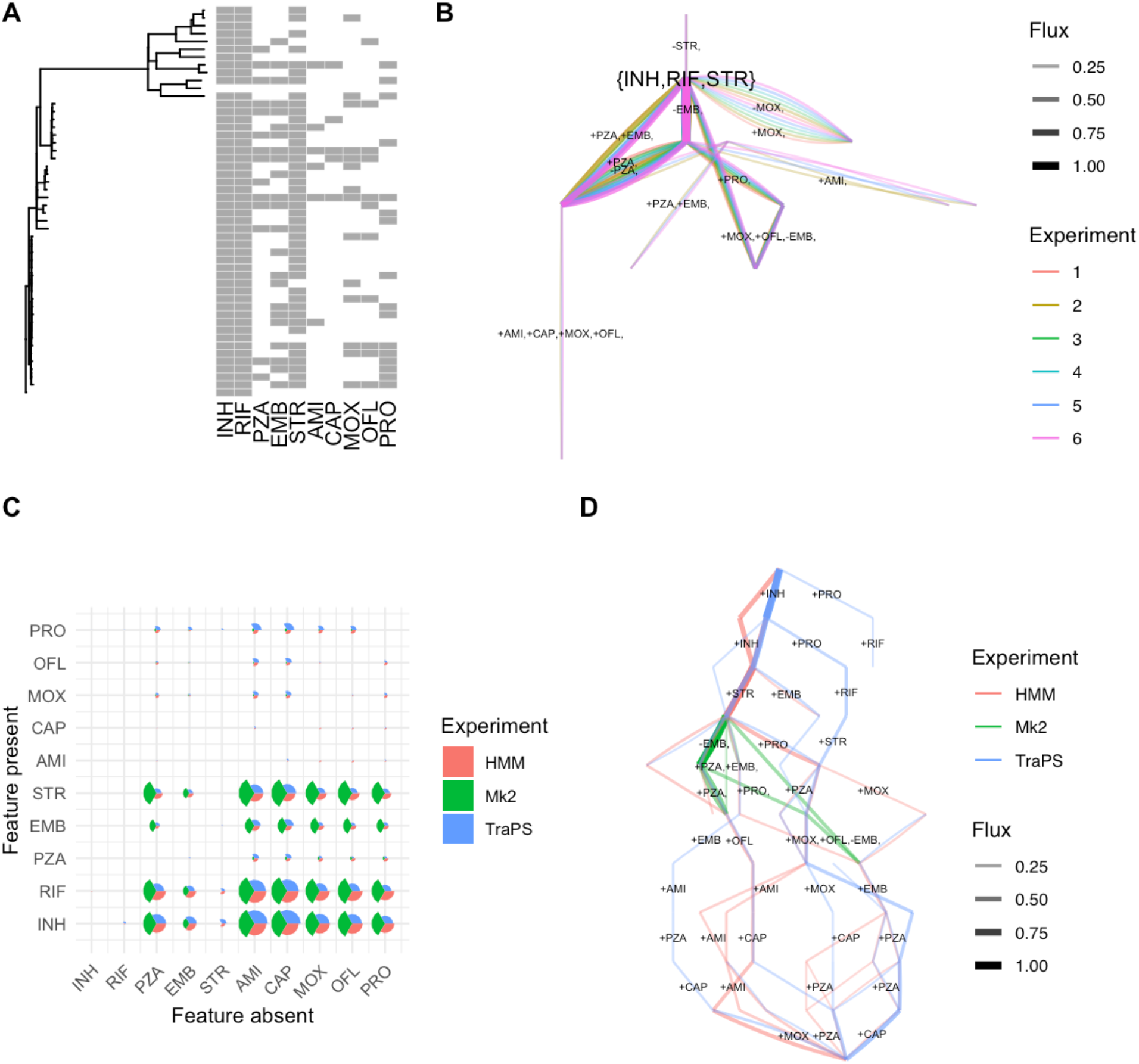
Inference and EvAM comparison for tuberculosis drug resistance data. (A) A subset of the data from (Casali et al., 2014), involving phylogenetically-linked tuberculosis isolates and patterns of resistance presence/absence against 10 drugs (label names given in Supp. Table 2). (B) Most common transitions in the inferred HyperMk2 transition network from these data, across 6 repeats of the algorithm, showing general consistency in the structure of reduced transition networks and their dynamics. (C) Relative ordering of pairs of features in HyperMk2 and other EvAM methods. Each segment’s size gives the probability that the “Before” feature is present when the “After” feature is acquired. This approach is “Mk2”; inferences using the irreversible approaches from HyperHMM “HMM” (Moen & Johnston, 2023) and HyperTraPS “TraPS” (Aga et al., 2024) are also shown. (D) Overlaid transition networks inferred from HyperMk2, HyperHMM, and HyperTraPS.

**Figure 4.**
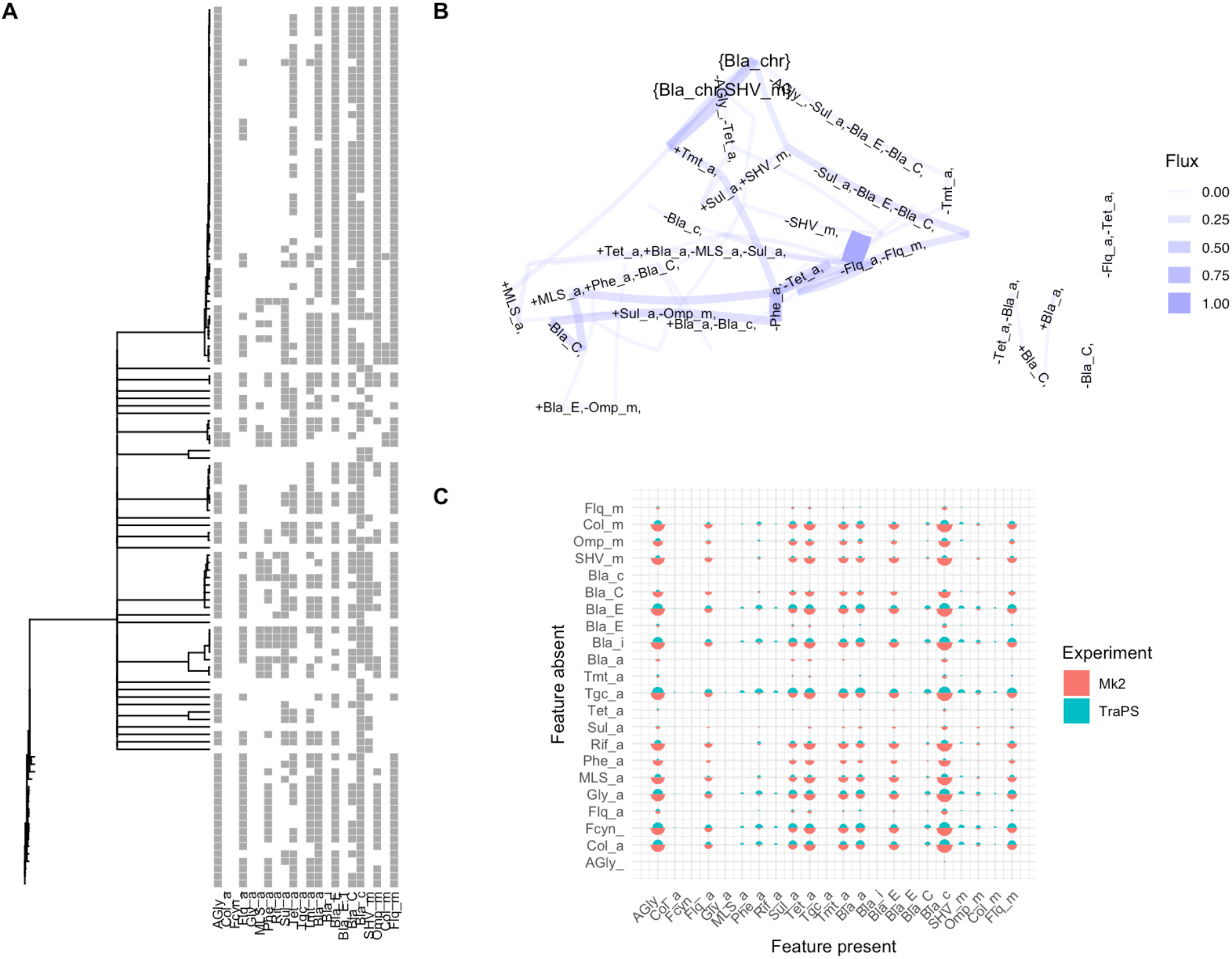
Inference for *Klebsiella pneumoniae* anti-microbial resistance (KpAMR) data. **(A)** Example KpAMR dataset from Romanian samples from PathogenWatch with features assigned using Kleborate (Argimón et al., 2021; ^L^am et al., 2021), from a global survey of KpAMR (Aga et al., 2025). ^L^abel explanations given in Supp. Table 2. **(B)** Most common transitions from the inferred transition graph from these data. The states involving only chromosomal resistance to beta-lactams (*Bla_chr*) and sulfhydryl variable mutations (*SHV_m*) are highlighted; these reflect common natural states of Kp. **(C)** Ordering matrix (as in Fig. 3C) showing the probability (segment size) with which features (“Before”) are likely present when an “After” feature is acquired. Orderings from HyperMk2 are compared to those from HyperTraPS in (Aga et al., 2025). The discrepancies are predominantly in *Bla_E, Bla_i, Tgc_a, Gly_a*, and *Fcyn_* features; these are features that do not appear in the Romanian data. *Col_m* and *Col_a* features are very rare in the dataset.

Fig. 3D compares the inferred transition networks from HyperTraPS, HyperHMM, and HyperMk2. HyperMk2 does not enforce evolution from a 000… precursor state (unlike the other approaches) and so has most probability flux from an {*INH, RIF, STR*} precursor state (the rare instances of absence of these features in Fig. 3A can be accounted for by losses in HyperMk2’s reversible picture). Likewise, HyperMk2 does not force walkers to a putative 111… state. The majority of probability flux from the HyperMk2 network is therefore present at those intermediate levels of the network reflecting the distribution of features in the dataset, where it agrees with the transition structure from the other approaches.

As before, we can compare the fitted model to a null model of independent character evolution to seek evidence for coupling between features. Here, the lowest AIC across the ensemble of reduced state spaces considered is 427.9, against an AIC of 455.2 for the independent case. However, unlike the bilinear case, it is not immediately clear from the structure of the data alone where the interactions driving this improvement may be. To characterise this coevolution, we search across pairs of features for evidence of co-dependence, by comparing an independent null model to a model allowing coupling between states (see Methods) (Boyko & Beaulieu, 2021a; Johnston & Diaz-Uriarte, 2025). This approach does not consider all pairwise interactions together as in other EvAM methods, but identifies the pairs for which strongest evidence for interaction exists. In this case, for example, ΔAIC is -11.2 for coupling *MOX* and *OFL* (the strong co-occurrence of which is visible in Fig. 3A). Examining the fitted model structure, we find that their acquisition is inferred to be tightly correlated and dependent on several other features being acquired (Fig. 3B,D). Similar, but weaker, positive influences are observed between *EMB* and *PZA*, and *AMI* and *CAP*. Indeed, interactions of *OFL* promoting *MOX* and *AMI* promoting *CAP* have previously been inferred from a larger dataset by HyperTraPS-CT (Aga et al., 2024), and reflect known mechanistic similarities (fluoroquinolone and aminoglycoside-peptide families respectively).

We also applied HyperMk2 to data on AMR in *Klebsiella pneumoniae* (KpAMR). Here, presence and absence of 22 KpAMR characters (Supp. Table 2) are inferred from genomes, and LINcodes, a coarse-grained measure of relatedness, are used to estimate phylogenetic connections between genomes (Aga et al., 2025; Hennart et al., 2022; Marakeby et al., 2014). Unlike the largely chromosomal changes in TB AMR, KpAMR characters are often gained and lost through horizontal gene transfer.

Reversibility is therefore a natural aspect of KpAMR evolution. While existing work has shown that useful information about reversible evolution can be gained from irreversible EvAM methods (Johnston, 2026), HyperMk2’s full support for reversibility allows a detailed analysis of this claim for the first time.

Fig. 4A shows a representative dataset from Romanian KpAMR samples. The reduced state space from Algorithm 1 is shown in Fig. 4B, and already reflects known biology: chromosomal features bestowing beta-lactam resistance (*Bla_chr*) and SHV mutation (*SHV_m*) are often constitutively present in Kp. Fig. 4C compares ordering matrices for the (reversible) HyperMk2 model and the (irreversible) HyperTraPS model used in (Aga et al., 2025). Agreement is observed across all characters, with aminoglycoside resistance (*AGly_*) appearing relatively early, tetracycline (*Tet_a*) and fluoroquinone (*Flq_m*) resistance at intermediate stages, and resistance to late-line drugs like colistin (*Col_m*) inferred as late acquisitions. There are several instances where the probability of character Y being absent when feature X is present is inferred to be higher in HyperTraPS than in HyperMk2, predominantly for X = *Bla_E, Bla_i, Tgc_a, Gly_a*, and *Fcyn_* features. These discrepancies arise because HyperTraPS extrapolates evolutionary trajectories to a final state of all features acquired, and it is in these extrapolated pathways that these relations arise; those KpAMR features are absent in this Romanian dataset and HyperMk2 therefore does not consider transitions involving their acquisition.

A comparison between a null model of independent character evolution and this model shows strong support for interactions between features (null model AIC 3223; HyperMk2 model AIC 1607). To identify interactions driving this difference, we did the same pairwise comparison of independent against paired models for character evolution (see Methods) to identify related pairs, and further organised by the magnitude of a Pearson residual for (uncorrected) correlation (Supp. Fig. 5). A collection of positive interactions are inferred through these criteria. Notably, these include *Tmt* and *Sul*, reflecting trimethoprim-sulfonamides, a common clinical drug combination, which is also identified as synergistic in nearly a quarter of country-specific datasets in (Aga et al., 2024)). Other identified pairings include *Tmt*/*Flq, Phe*/*MLS*, and *Phe*/*Flq*, which may correspond to joint carriage on resistance cassettes and/or joint resistance through efflux pump dynamics.

## Discussion

Previous work has established a hypercubic instance of the Mk model as a useful picture for considering the coupled acquisition of multiple features through related lineages in evolution. Dependency relationships between multiple characters may be expected in many evolutionary circumstances, from shared mechanisms and collateral sensitivity in drug resistance (Baquero et al., 2021; Pál et al., 2015; Renz, Dauda, et al., 2025), competing (mutually repressing) pathways in metabolic evolution (Glastad & Johnston, 2025; Williams et al., 2013), and the evo-devo of multiple characters (Boyko & Beaulieu, 2021b). Previous EvAM approaches for phylogenetically-linked data have assumed irreversibility and perfect observation (and some make strict consequent assumptions about ancestral state reconstruction) (Aga et al., 2024; Luo et al., 2023; Moen & Johnston, 2023; Renz, Brun, et al., 2025). The hypercubic Mk model picture has the notable advantage of supporting reversibility and consequent flexibility in ancestral state reconstruction for phylogenetically embedded data, while retaining support for uncertain observations and arbitrary positive and negative interactions between characters (Johnston & Diaz-Uriarte, 2025). But this flexibility previously came at the cost of computational complexity, with aggressive exponential scaling of runtime (and poor guarantees on convergence) with number of features *L*, making a wide range of interesting problems intractable in practise. An optimistic perspective on this article would be that we retain that flexibility while taking large steps to overcoming the problems associated with poor scaling. Empirically, for the time *t* in seconds to fit a model involving randomly-simulated (unstructured) data with *s* states in the reduced space and *n* observations, on a single laptop core, we observe *t* ≈ 1.52 × 10^-7^ *s*^4.44^ *n*^1.14^. An unstructured problem involving, for example, 1000 observations over 100 states would be predicted to take roughly 86 hours to fit. Highly structured problems (like the linear pathway) scale much less dramatically (sublinearly) with both *s* and *n*, and are fitted in orders of magnitude less time.

But we have not achieved a free lunch. Our approach for reducing the size of the state space removes a large number of evolutionary possibilities from consideration in the inference process. Two classes of possible problem can result: the unbiased reduction of potentially important dynamics, and any systematic biases in the way we construct our reduced space size.

The first class of problem is in some sense a feature, not a bug, of our approach. We explicitly invoke a minimum-evolution picture, asserting that less complex evolutionary pathways are more likely. This is true in the sense that more extended cycles of repeated gain and loss cycles are less likely for any transition probabilities under unity, in the absence of further information. But it may nevertheless be the case that true evolutionary history involved many multiple repeated gains and losses of reversible characters, and our approach has no means of capturing this. Although cautiously used in some EvAM applications (Dauda et al., 2025), it is classically known that minimum evolution pictures may be flawed for reconstructing phylogeny in even simple cases (Felsenstein, 1978); here our use of this picture is instead to consider a plausible simplest state-space model that is capable of accounting for observed diversity in characters.

The main concern regarding the second class of problem, systematic biases, is probably the greedy way in which our algorithm forms putative sets of states for each node from local information alone. Without an exhaustive search over the tree, it is possible that a judicious choice of ancestral state for some node may allow a more parsimonious description of the dynamics of several lineages. The establishment of most-parsimonious evolutionary pathway sets is NP-hard in several cases (Giannakis et al., 2024), so a perfect solution is likely to remain elusive. However, the interpretation of outputs from our approach should involve a consideration of the particular set of ancestral states involved and whether they are sufficiently representative of a parsimonious evolutionary picture. It is encouraging that differently sampled instances of the algorithm remain consistent in the transition networks and summary dynamics inferred (Fig. 3B, Supp. Figs. 2, 4), but further testing of this consistency will increase the robustness of this approach. The agreement of available dynamic summaries between irreversible EvAMs and HyperMk2 also paints a positive picture about the previous use of irreversible methods (Johnston, 2026).

A further potential advantage of this method is its interoperability. As the algorithm fundamentally provides a Q-matrix, set of states, and tree for the Mk model fit, the wealth of available methods for fitting and analysing the Mk model can be applied (Beaulieu et al., 2022; Louca & Doebeli, 2018; Revell, 2024) and at the same time the collection of methods for analysing EvAM outputs can be applied downstream (Aga et al., 2024; Diaz-Uriarte & Herrera-Nieto, 2022). Model comparison (for example, via AIC, as used in this report) can readily be used to compare different cases and null models (Boyko & Beaulieu, 2021b) and positive and negative controls for particular computational experiments can readily be designed (Aga et al., 2024). One particular addition in this report is the identification of specific interactions between features in reversible, phylogenetic EvAM: previously, HyperMk provided a flexible state space framework but did not explicitly identify particular pairwise interactions as with other EvAM approaches (Aga et al., 2024; Hjelm et al., 2006; Schill et al., 2020). We hope that this flexibility helps open a new set of research domains for the growing field of EvAM.

## Acknowledgements

This work was supported by the Trond Mohn Foundation (project HyperEvol under grant agreement No. TMS2021TMT09 to I.G.J.) through the Centre for Antimicrobial Resistance in Western Norway (CAMRIA) (TMS2020TMT11). This project has received funding from the European Research Council (ERC) under the European Union’s Horizon 2020 research and innovation program (grant agreement No. 805046 [EvoConBiO] to I.G.J). This work was supported by grant PID2024-156888OB-I00 funded by MICIU /AEI/10.13039/501100011033 / FEDER, EU to R.D-U.

## Supplementary Information

Supplementary Text 1. **Example of progress through Algorithm 1 for the illustration in Fig. 1**. † denotes the sampling choice made differently (001) for the case in Supp. Fig. 1.

**Input:** Tree ((t1,t2)n7,(t3,t4)n6)n5; tip states t1 = 100, t2 = 101, t3 = 010, t4 = 001

A. *Construct set of possible labellings* (Fig. 1(i)-(iii))

[Start]

Node n6 has labelled daughters t3, t4

Both daughters are tips

Assign 010 ⊕001 = {000, 001, 010, 011} to node n6

Node n7 has labelled daughters t1, t2

Both daughters are tips

Assign 100 ⊕101 = {100, 101} to node n7

Node n5 has labelled daughters n6, n7

Both daughters are not tips

Let {*D, W*_1_, *W*_2_}=*D*_1_ ⊕ *D*_2_ = {{000, 100, 001, 101}, {{000}, {000}, {001}, {001}}, {{100}, {100}, {101}, {101}}

Assign {000, 100, 001, 101} to node n5 and record {*W*_1_, *W*_2_}

[All vertices are now labelled]

B. *Sample a locally consistent possible labelling* (Fig. 1(iv))

Call Recursive-choice(n5, {000, 100, 001, 101})

Randomly sample 100 for node n5 †

Node n5 has daughters n6, n7

Call Recursive-choice(n6, {000})

(Randomly) sample 000 for node n6

Node n6 has no daughters without assigned states

Call Recursive-choice(n7, {100})

(Randomly) sample 100 for node n7

Node n7 has no daughters without assigned states

[Done]

**Output:** (Fig. 1(v))

[From n5 → n6:] 100 → 000

[From n5 → n7: no transition]

[From n6 → t3:] 000 → 010

[From n6 → t4:] 000 → 001

[From n7 → t1: no transition]

[From n7 → t2:] 100 → 101

This set of states is then indexed to form a contiguous set of natural numbers (000 = 1, 001 = 2, 010 = 3, 100 = 4, 101 = 5), and the Q-matrix for the Mk model fit therefore is constrained to be zero everywhere except elements corresponding to 4 → 1, 1 → 3, 1 → 2, 5 → 4. The Mk model is then fit using the original tree, this Q-matrix constraint, and tip states from the index set t1 = 4, t2 = 5, t3 = 3, t4 = 2.

Supplementary Text 2. **Algorithm 2 – Reducing state space for a hypercubic Mk model (non-recursive)**

**Input:** a phylogenetic tree with binary string observations of length *L* at each tip Label all tips with their binary-string observations

While unlabelled nodes remain

Loop over nodes

If this node has labelled daughters *D*_1_, D_2_

If both daughters are tips

Assign *D*_1_ ⊕*D*_2_ to this node

Otherwise

Let {*D, W*_1_, W_2_}=D_1_ ⊕ *D*_2_

Assign set {*D*} to this node and {*W*_1_, W_2_} to the two daughters ^*^

End if

End if

End loop

End while

For all nodes with state sets consisting of more than one element Randomly choose an element from the state set list

Construct state space consisting of ancestor-node transitions for each branch on the chosen tree

**Output:** set of states and transitions between them, to constrain the Mk model Q-matrix

^*^ To enforce 000… as the root state, if this node is the root, replace {*D*} with 000… and {*W*_1_, W_2_} with witness sets from descendent sets giving minimum Hamming distances to 000… .

**Supplementary Figure 1.**
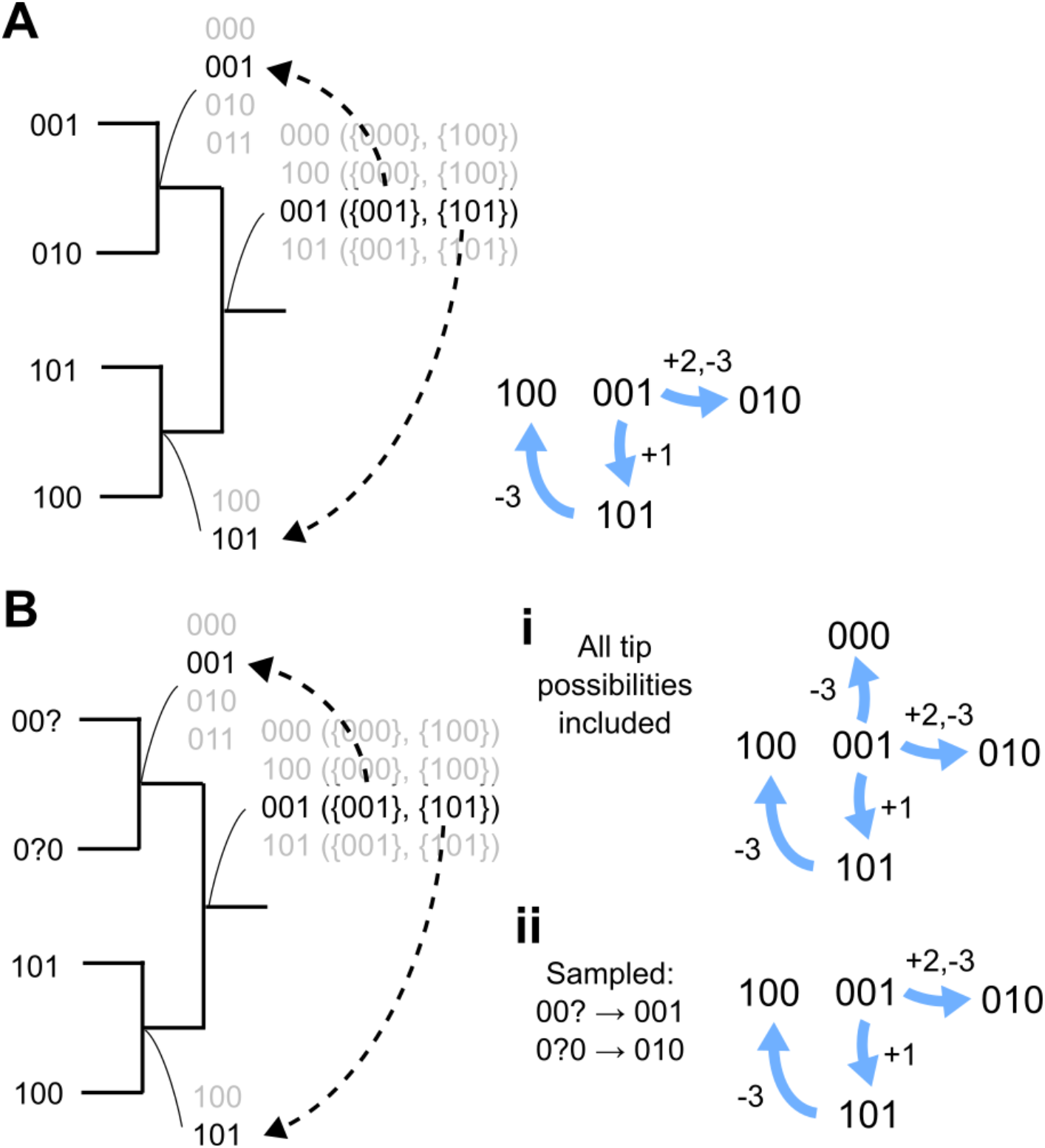
Alternative instances of Algorithm 1. **(A)** The data and state construction are the same, but now state 001 is sampled for the root (Supp. Text 1). One resultant transition goes from 001 to 010, involving two changes (a loss and a gain). **(B)** The tip data now includes uncertainty (?) in observations. Each uncertain feature is expanded out to form the ancestral state set. To sample a given instance, we can (i) retain all tip states (and transitions to them) compatible with the uncertain observations; or (ii) sample a particular compatible instance of tip states per instance of the algorithm.

**Supplementary Figure 2.**
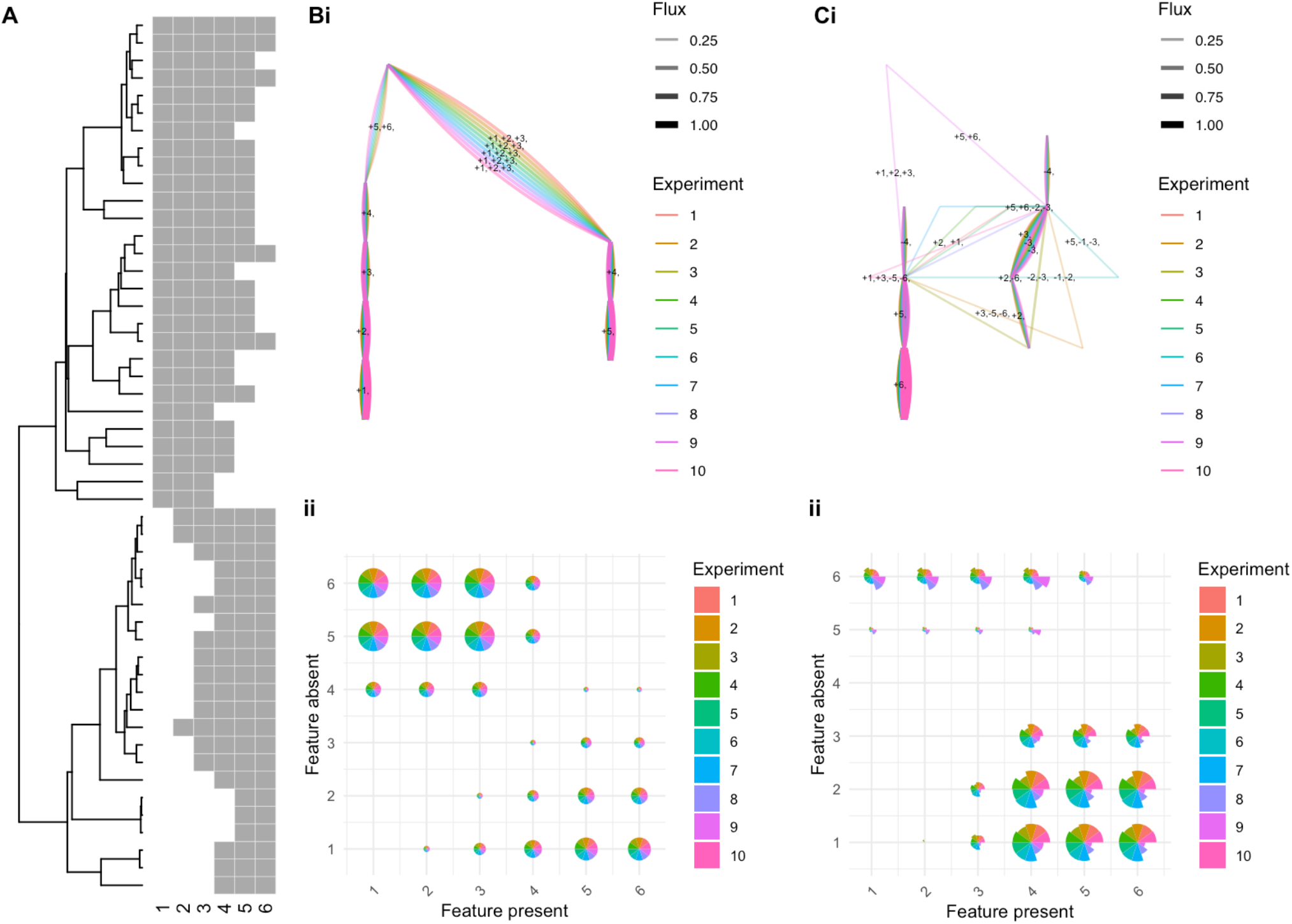
Consistency of algorithm solutions across sampled instances. (A) Data from the synthetic case study in Fig. 2. (B) Irreversible algorithm: (i) inferred transition networks and (ii) ordering matrices from 10 instances of the algorithm (different colours). (C) Reversible algorithm: (i) inferred transition networks and (ii) ordering matrices from 10 instances of the algorithm (different colours).

**Supplementary Figure 3.**
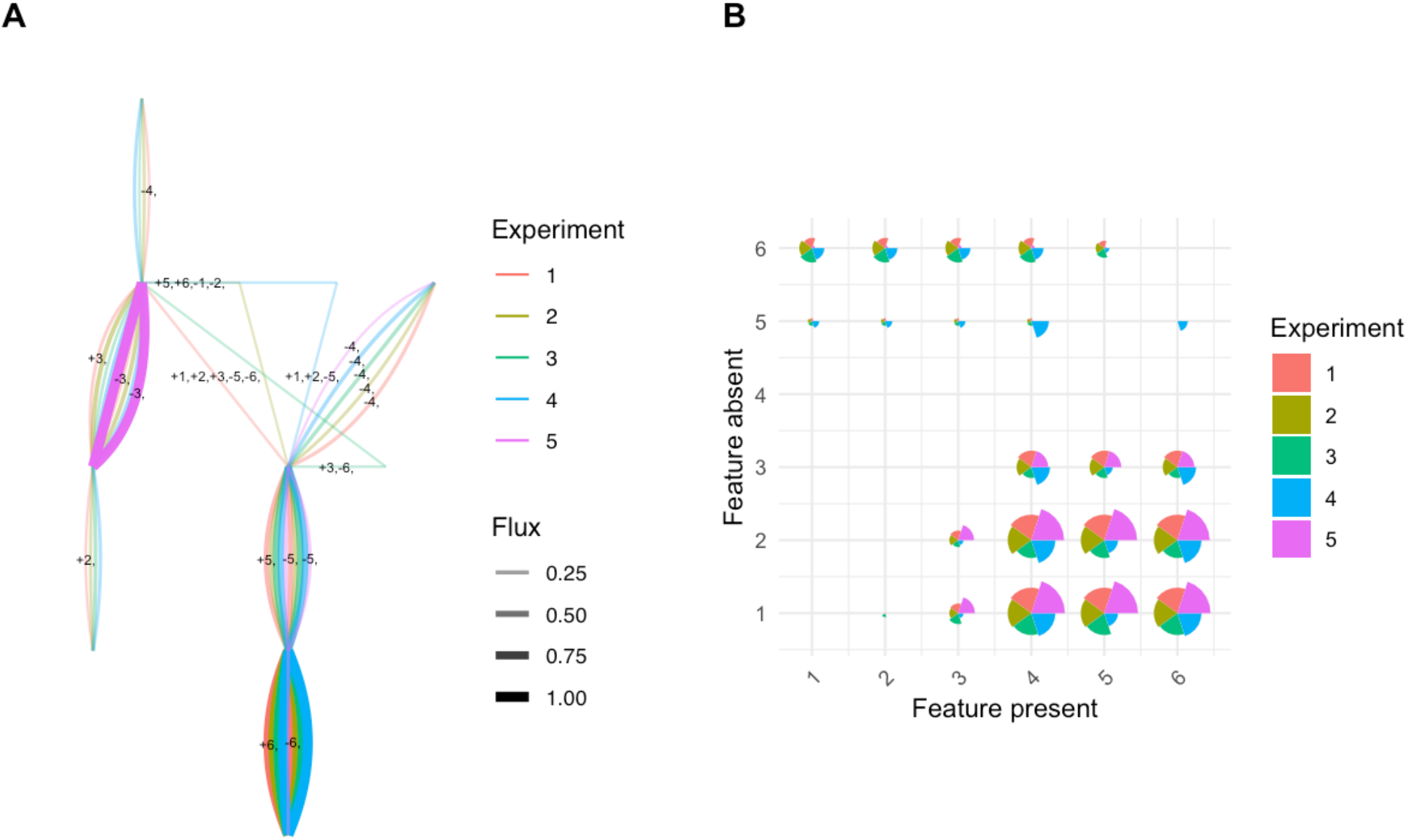
Performance under different levels of uncertainty. (A) Inferred transition networks and (B) ordering matrices from the algorithm applied to the data in Fig. 2 with increasing synthetic occlusion of observations: 1, 0%; 2, 10%; 3, 20%; 4, 30%, 5, 40% occluded.

**Supplementary Figure 4.**
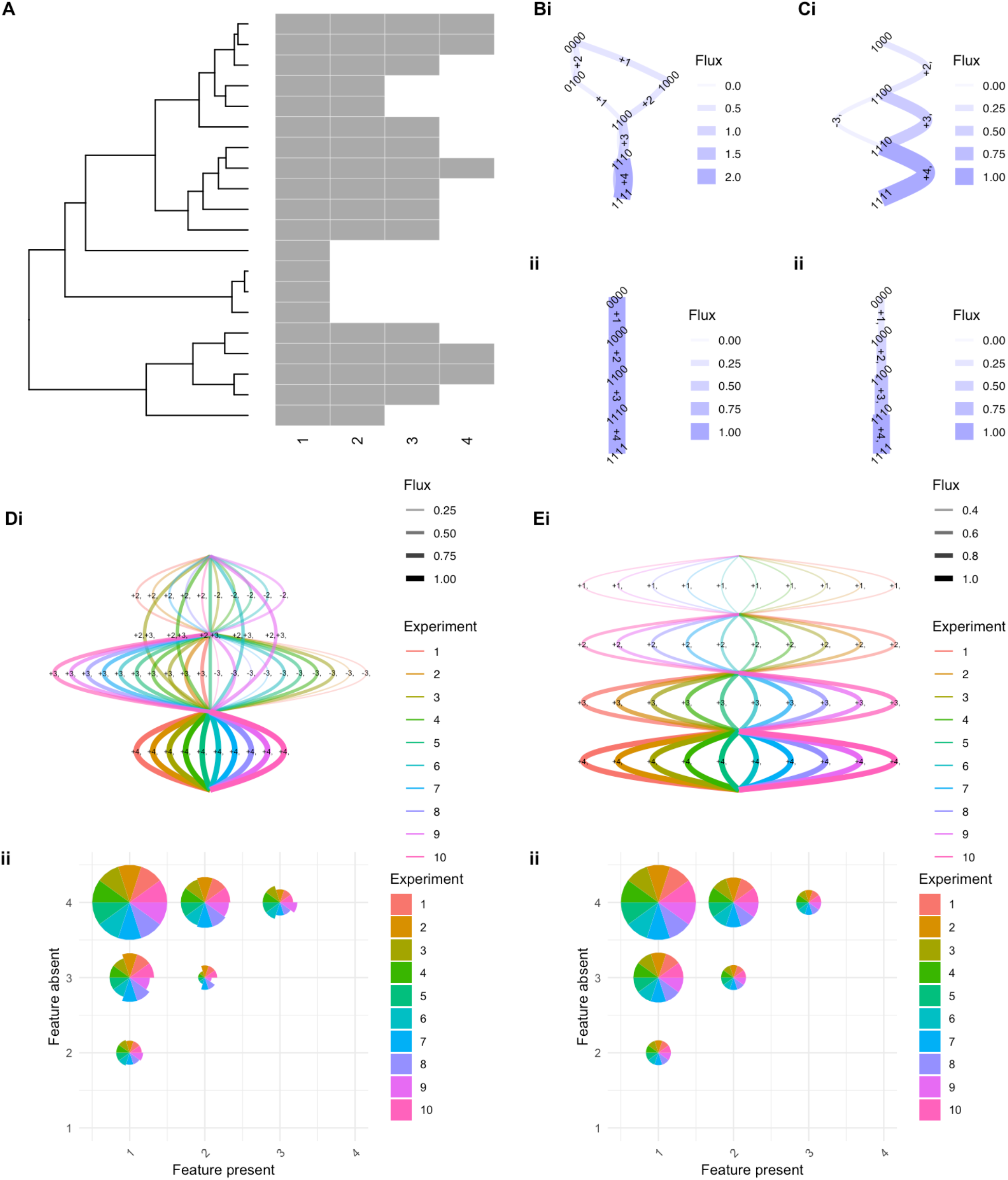
Application of the method to a linear test dataset. (A) A simulated dataset involving unidirectional, monotonic accumulation from features 1 to 4. (B) HyperMk (C) HyperMk2 fits assuming (i) reversible (ii) irreversible pictures. (D) Reversible HyperMk2 (i) transition networks and (ii) ordering matrices across 10 instances of the algorithm (different colours). (E) Irreversible HyperMk2 (i) transition networks and (ii) ordering matrices across 10 instances of the algorithm (different colours).

**Supplementary Figure 5.**
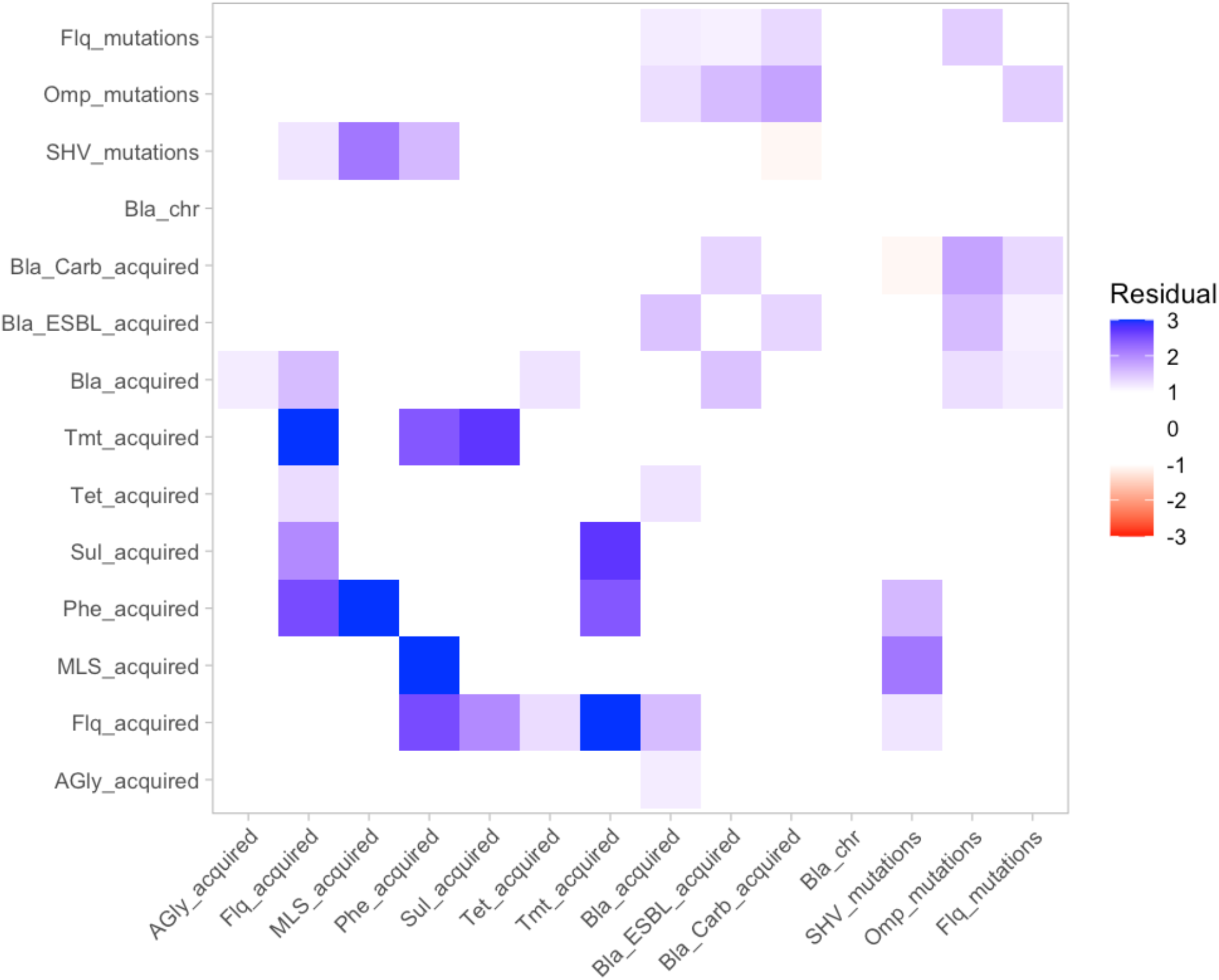
Interactions in the Kp AMR dataset. For each pair of characters we first construct a standardised Pearson residual 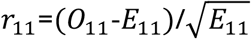 where *E*_11_=(*n*_.1_+*n*_11_)(*n*_1._+*n*_11_)/*N*, and pairs of numbers refer to particular presence/absence combinations. This value is computed without accounting for phylogeny. We then compute the ΔAIC measure from Methods, using the phylogenetic information to compare support for an independent-characters model against one of paired evolution. For pairs with ΔAIC < -10, we plot the value of the Pearson residual, highlighting those values at or exceeding 2 in magnitude.

**Supplementary Table 1.** Functions in the HyperMk2 R package.

| Function name | Description | Arguments | Output |
| --- | --- | --- | --- |
| build_states | Fitch-like algorithm to build possible minimum-evolution state sets over a tree | Phylogenetic tree<br>Matrix of binary observations (one row per tree tip) | A nested list. Each element corresponds to a tip or node in the tree. Each subelement corresponds to a possible state at that tip, along with a matrix describing the indices of "witness" states at the descendant vertices that correspond to the given state at this vertex. |
| cheap_transition_set | Build a cheap approximation to reduced state space, avoiding Fitch-like computation | Phylogenetic tree<br>Matrix of binary observations (one row per tip) | A dataframe with transitions between (decimal) states in the reduced space. |
| hypermk2 | Use HyperMk2 to fit a reversible evolutionary accumulation model | Phylogenetic tree<br>Matrix of binary observations (one row per tip)<br>Reversible model (Boolean, default TRUE)<br>Whether to force the root to have state 000... (Boolean, default FALSE)<br>Whether to compare a null model of independent features (Boolean, default FALSE)<br>Whether to use cheap state space algorithm (Algorithm 2) (Boolean, default FALSE) | A named list containing the fitted Mk model object, inferred fluxes between states, the number of features, set of transitions in the reduced space, and feature names |
| hypermk2_independent | Get statistics of a null model fit of independent features | As hypermk2 | A named list containing total and feature-specific likelihoods and AICs |
| phylo_correlations | Get correlation statistics of pairs vs independent features | Phylogenetic tree<br>Matrix of binary observations (one row per tip) | A named list containing ΔAICs and AICs for independent and coupled model fits |
| sample_states | Sample a single minimum-evolution instance from the Fitch-like algorithm output | Phylogenetic tree<br>Nested list describing node labels (output from build_states) | A named list containing (a) a list of binary vectors, with each vector corresponding to the sampled state at that vertex in the tree; (b) a dataframe of From-To transitions on the tree (in decimal representation) |

**Supplementary Table 2.**
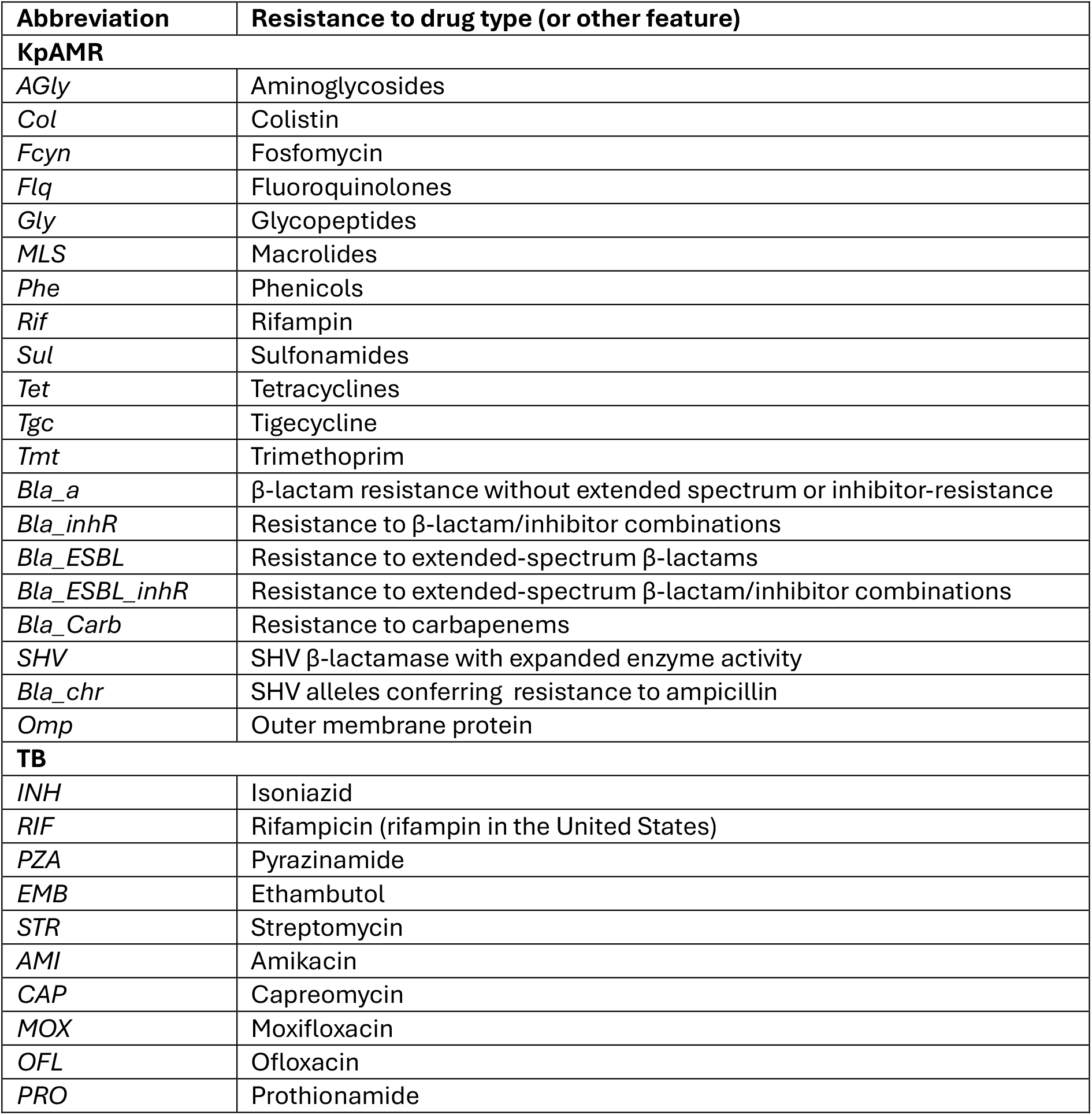
AMR character names. More details available via (TB) (Aga et al., 2024; Casali et al., 2014); (KpAMR) Kleborate documentation at https://github.com/klebgenomics/Kleborate and (Aga et al., 2025).

